# Oxysterol Alterations in SOD1G93A ALS Rats: 25-Hydroxycholesterol and LPS-Binding Protein in Disease Progression

**DOI:** 10.1101/2025.05.05.650333

**Authors:** Rodrigo Santiago Lima, Rosangela Silva Santos, Adriano Britto Chaves-Filho, Isis Ferraz Libório Trzan, Alex Inague, Rodrigo Lucas de Faria, Lucas Souza Dantas, Isabella Fernanda Dantas Pinto, Marisa H.G. Medeiros, Alexandre Alarcon Steiner, Sayuri Miyamoto

**Author notes:** Corresponding author: Instituto de Química, Departamento de Bioquímica, Universidade de São Paulo, CP 26077, CEP 05513-970, São Paulo, Brazil.

## Abstract

**Background:** Disruptions in cholesterol and oxysterol metabolism, along with neuroinflammation, are linked to amyotrophic lateral sclerosis (ALS), though the underlying mechanisms remain unclear. Given evidence of increased intestinal permeability in ALS, we investigated its link to neuroinflammation and oxysterol alterations in SOD1G93A rats.

**Methods:** Oxysterols were quantified in plasma and spinal cord from presymptomatic and symptomatic SOD1G93A rats and age-matched controls via ultra-high performance liquid chromatography coupled with high-resolution mass spectrometry. Circulating LBP, a marker of intestinal permeability, was quantified via ELISA.

**Results:** Oxysterols involved in bile acid biosynthesis - 7α-hydroxycholesterol, 27-hydroxycholesterol (27-OH), and 3β-hydroxycholestenoic acid - were increased in the plasma of symptomatic rats. The neuronal oxysterol 24(S)-hydroxycholesterol (24(S)-OH) decreased in the spinal cord but increased in the plasma. In contrast, 27-OH and 25-hydroxycholesterol (25-OH) levels were elevated in both plasma and spinal cord, with 25-OH rising during the presymptomatic stage. Presymptomatic animals also exhibited elevated LBP levels, which strongly correlated with spinal cord 25-OH levels, suggesting a link between systemic inflammation and neuroinflammation in ALS.

**Conclusion:** Oxysterol alterations in plasma and spinal cord suggest compromised blood‒spinal cord barrier integrity and early neuroinflammation. Elevated LBP levels indicate increased intestinal permeability and circulating LPS as contributors to neuroinflammation and neurodegeneration. These findings highlight 25-OH and LBP as markers and mediators of gut‒brain axis interactions in ALS pathogenesis, particularly in the presymptomatic phase.

## INTRODUCTION

The brain, a cholesterol-rich organ, contains approximately 25% of the body’s total cholesterol [1]. Cholesterol is essential for cell membrane integrity, myelin sheath maintenance, and various physiological processes [2]. The blood‒ spinal cord barrier (BSCB), similar to the blood‒brain barrier (BBB), consists of tightly packed endothelial cells, a basal membrane, pericytes, and astrocyte endfeet [3]. Both barriers protect the central nervous system (CNS) from fluctuations in blood cholesterol levels by isolating it from systemic circulation [1]. To maintain cholesterol homeostasis, the CNS metabolizes cholesterol into oxysterols, which regulate cholesterol synthesis, uptake, and excretion [2, 4–6].

Oxysterols are formed through both enzymatic and nonenzymatic oxidation of cholesterol. While the cytochrome P450 (CYP450) family of enzymes plays a central role in oxysterol production [7, 8], other enzymes, such as cholesterol 25-hydroxylase (CH25H) [9] and HSD3B7 [10], are also involved. In the CNS, CYP46A1 is the key enzyme responsible for maintaining cholesterol homeostasis. Primarily expressed in the brain [5], more specifically in neurons [11], CYP46A1 catalyzes the conversion of cholesterol into 24(S)-hydroxycholesterol, which can cross the BBB/BSCB, providing an essential route for excess cholesterol excretion from the brain [12].

Beyond cholesterol regulation, oxysterols serve as precursors for bile acid and steroid hormone biosynthesis [10, 13]. They also act as ligands for nuclear receptors such as liver X receptors (LXR) [14] and G protein-coupled receptor 183 (GPR183 or EBI2) and regulate the INSIG-SCAP-SREBP complex. LXRs control the expression of genes involved in cholesterol trafficking, including ABCA1, ApoE and IDOL [15], while the INSIG-SCAP-SREBP complex regulates genes responsible for cholesterol synthesis and uptake [16–18].

Alterations in oxysterol levels have been reported in several neurodegenerative diseases, including Alzheimer’s disease [19–26], cerebrotendinous xanthomatosis, spastic hereditary paraplegia [27], Huntington’s disease [28–30], Niemann‒Pick disease [31, 32], Smith‒Lemli‒ Opitz syndrome [33] and amyotrophic lateral sclerosis (ALS) [34–38]. Investigating the mechanisms underlying oxysterol dysregulation in neurodegeneration may provide insights into new therapeutic strategies or ways to slow disease progression.

In ALS, some hallmarks include disruption of the BSCB, as evidenced by reduced expression of tight junction proteins [39, 40] and collagen perturbations [41–44]; increased intestinal permeability, as indicated by decreased tight junction proteins between epithelial cells; and elevated plasma levels of LBP (LPS-binding protein) [45–47]. Neuroinflammation is another hallmark, as evidenced by activated microglia and astrocytes [48–52] and immune cell infiltration in the spinal cord [53–56]. The idea that these pathological characteristics may somehow communicate with each other and ultimately lead to neurodegenerative processes is a possibility that cannot be overlooked.

In this study, we performed precise quantification of oxysterols in the blood plasma and spinal cord of a SOD1G93A ALS rat model using ultra-high performance liquid chromatography coupled with high-resolution mass spectrometry (UHPLC‒QTOF‒MS). Oxysterol alterations in the spinal cord and plasma indicate that BSCB disruption occurs at late stages of ALS and reveal evidence of neuroinflammation even at asymptomatic stages. Additionally, we quantified plasma LBP as a marker of intestinal permeability [57, 58] and reported that LBP levels were elevated in presymptomatic SOD1G93A rats and correlated positively with 25-hydroxycholesterol (25-OH) levels in the spinal cord. This correlation suggests a link between systemic inflammation, intestinal permeability and neuroinflammation in ALS.

## MATERIALS AND METHODS

### Animals and sample preparation

The study followed the ethical guidelines of the National Animal Experimentation Control Council (CONCEA) and was approved by the Ethics Committee on Animal Use of the University of São Paulo (CEUA 41/2016). Male Sprague Dawley rats overexpressing the SOD1G93A gene, obtained from Taconic, were housed under controlled temperature conditions with a 12 hour light/dark cycle. Water and food were provided ad libitum. Rats were classified as asymptomatic until 90 days of age and symptomatic at 120 days, on the basis of criteria from Matsumoto [59]. Animals showing muscle paralysis or 15% body weight loss were euthanized to minimize suffering. Prior to sample collection, the rats were fasted for 6 hours and anesthetized with isoflurane. Asymptomatic rats (SOD1G93A 90d; n=9) and age-matched wild-type controls (n=10 for plasma and n=9 for spinal cord samples) were sacrificed at 89 ± 3 days of age. Symptomatic rats (SOD1G93A 120d; n=10 for plasma and n=9 for spinal cord samples) and age-matched controls (n=10 for plasma and n=9 for spinal cord samples) were sacrificed at 120 ± 6 days of age. Blood was collected by cardiac puncture into a tube containing heparin, and the plasma was separated by centrifugation at 2000 × g for 10 minutes at 4 °C. Spinal cords were frozen immediately after dissection and macerated using a mortar and pestle with liquid nitrogen to prevent thawing. Approximately 200 mg of tissue was transferred to Eppendorf tubes, followed by the addition of 5 mM phosphate buffer (pH 7.4) containing 10 µM deferoxamine mesylate salt (Sigma‒Aldrich). The tissue was homogenized via a tissue homogenizer to achieve a final concentration of 200 mg/mL. Both plasma and spinal cord homogenate samples were stored in a −80 °C freezer until further analysis.

### Oxysterol standards

For the analysis, we used a “mix of oxysterols” containing both commercially available and synthesized compounds (**Table S1**). The commercially obtained standards were purchased from Avanti Polar Lipids and included 7-ketocholesterol (code: 700015P), 7α-hydroxycholesterol (code: 700034P), 5β,6β-epoxycholesterol (code: 700033P), 25-hydroxycholesterol (code: 700019P), 24(S)-hydroxycholesterol (code: 700061) and 27-hydroxycholesterol (code: 700021). Additional standards, including 4β-hydroxycholesterol (Item No. 19518), 22(R)-hydroxycholesterol (Item No. 89355), and 3β-hydroxy-5-cholestenoic acid (Item No. 21859), were acquired from Cayman Chemical Company. Standards for 5α-hydroxycholesterol, 6β-hydroxycholesterol, 3β-hydroxy-5-oxo-5,6-secocholestan-6-al (Secosterol A or CSec) and 3β-hydroxy-5β-hydroxy-B-norcholestane-6β-carboxyaldehyde (Secosterol B or ChAld) were synthesized in our laboratory as previously described [60]. The internal deuterated standards used in the analysis included d6 24-hydroxycholesterol (code: LM-4110), d7 7-ketocholesterol (code: LM-4107) and d7 7α-hydroxycholesterol (code: LM-4103) from Avanti Polar Lipids, as well as d6 cholesterol (code: 488577) from ISOTEC. All sterol solutions were prepared using HPLC-grade isopropanol.

### Oxysterol extraction

The oxysterols extraction method was based on protein precipitation from biological matrices via the use of an organic solvent [61, 62]. To each 2 mL Eppendorf tube, we added 100 µL of oxysterol internal standards (d6 24-OH, d7 7α-OH and d7 7-keto 0.2 µg/mL [0.5 µM] in IPA) and 50 µL (or 100 µL for the spinal cord homogenates) of d6 cholesterol (1 mg/mL). Subsequently, 200 µL of ethanol containing 10 µM BHT was added. Plasma (200 µL) or the spinal cord homogenate (100 µL) was then added dropwise, followed by vortexing and incubation on ice for 15 minutes. Two more additions of 400 µL of ethanol were subsequently made, followed by the final addition of 650 µL (700 µL in the case of the spinal cord samples). After each addition, the samples were vortexed and incubated on ice for 15 minutes. After protein precipitation, the samples were centrifuged at 10,000 × g for 10 minutes at 4 °C. The supernatant (1.6 mL) was transferred to a 2 mL vial and dried under nitrogen flow. The dried residue was subsequently resuspended in 80 µL of isopropanol.

### Oxysterol analytical method

Oxysterol analysis was performed using a UHPLC-QTOF-MS system consisting of ultra-high performance liquid chromatography (UHPLC Nexera, Shimdazu, Kyoto, Japan) coupled with a high-resolution quadrupole time-of-flight mass spectrometer (TripleTOF^®^ 6600, Sciex, Concord, US). A 4 µL sample was injected into a C18 column (Acquity UPLC BEH^®^, 100 mm × 2.1 mm i.d., 1.7 µm, Waters), with a flow rate of 0.35 mL/min and an oven temperature of 20 °C. For reversed-phase liquid chromatography, mobile phase A consisted of water/acetonitrile (1:1), and mobile phase B consisted of an acetonitrile/isopropanol (1:1) mixture, both of which contained 0.1% formic acid. Oxysterols separation was achieved via a 20 min gradient: 20% to 50% B over the first 0.5 min, 50 to 100% B from 0.5 to 9 min, 100% B from 9 to 16 min, 20% B from 16 to 16.5 min, and 20% B from 16.5 to 20 min.

Mass spectrometry analysis was conducted using an electrospray ionization (ESI) source in positive mode with the following parameters: curtain gas at 25 psi, nebulizer (GS1) and heater (GS2) gas at 55 psi, ion spray voltage at 5500 V, and ion source temperature at 550 °C. The mass-to-charge scan range was set from 200 to 1000 Da (MS). The following m/z values were monitored for the oxysterol analytes: 419.3470 ± 0.005 (ChAld and CSec), 401.3364 ± 0.005 (7-keto), 399.3258 ± 0.005 (3β-HCA), 385.3465 ± 0.005 (monohydroxysterols), 383.3258 ± 0.005 (ChAld and CSec), 369.3515 ± 0.005 (cholesterol) and 367.3359 ± 0.005 (monohydroxysterols) (**Figure S1**). For the deuterated internal standards, the monitored m/z values were 408.3803 ± 0.005 (d7 7-keto), 391.3838 ± 0.005 (d6 24-OH), 375.3985 ± 0.005 (d6 cholesterol), 374.3749 ± 0.005 (d7 7α-OH) and 373.3739 ± 0.005 (d6 24-OH).

### Oxysterol quantification

Oxysterol quantification was achieved using a calibration curve. The oxysterol mixture solution at an initial concentration of 2 µM was serially diluted to produce 8 calibration points. The calibration curve was analyzed in quintuplicate via the LC‒QTOF‒MS system. Each oxysterol peak area was integrated using the MultiQuant™ 3.0.3 software (AB Sciex), and parameters such as linearity (R^2^), coefficient of variation (CV%), accuracy and limit of quantification (LOQ) were calculated. A weighing factor of 1/x^2^ was applied to the curves based on data from the literature regarding guidelines in LC‒MS-based calibration curves [63]. The quintuplicate analysis of all 8 calibration curve points allowed us to obtain a reliable equation for each pair of analyte/internal standard (**Table S2**). The equations used to calculate the oxysterol concentration followed the linear regression model of *y = ax + b*, where *y* was the area ratio (analyte peak area/IS peak area) and *x* was the concentration injected.

### LBP quantification

Plasma LBP (lipopolysaccharide-binding protein) quantification was performed in duplicate via the Novus Biologicals Rat LBP ELISA Kit (NBP2-75370) following the manufacturer’s protocol. The plasma samples were diluted 1:10 to fit within the LBP standard curve range.

## RESULTS

### The oxysterol profile of SOD1G93A rats indicates BSCB disruption and inflammation

To investigate systemic changes in oxysterol levels, we quantified these compounds in the plasma of ALS SOD1G93A rats. Symptomatic ALS rats presented elevated plasma concentrations of 24(S)-OH (Fig. 1), a cholesterol metabolite primarily synthesized in the central nervous system. The increase in 24(S)-OH was significant when symptomatic SOD1G93A rats at 120 days (ALS120) were compared with both asymptomatic 90-day-old SOD1G93A rats (ALS90) and age-matched controls. The increase in 24(S)-OH suggests that disease progression is associated with either increased cholesterol metabolism in the central nervous system and/or increased BSCB permeability. Another notable finding was the elevated plasma concentration of 25-OH, an oxysterol typically associated with inflammatory responses. Compared with asymptomatic ALS rats (ALS90) and age-matched controls (WT90 and WT120), symptomatic SOD1G93A rats (ALS120) presented significantly higher 25-OH plasma levels (Fig. 1), indicating that the inflammatory process intensifies as the disease progresses.

**Figure 1.**
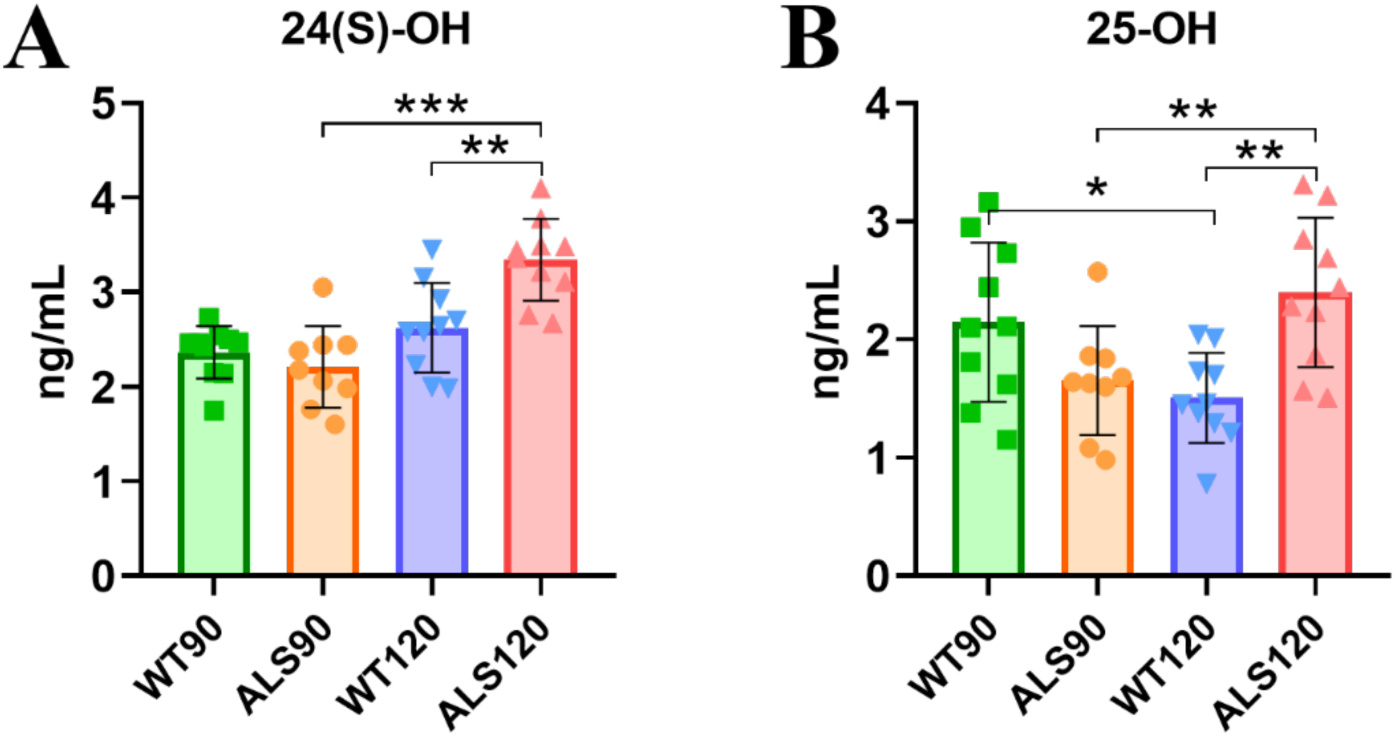
Increased plasma concentrations of 24(S)-OH (**A**) and 25-OH (**B**) in ALS symptomatic animals. Oxysterol levels were measured in the blood plasma of SOD1G93A rats and age-matched controls at the asymptomatic (89 ± 3 days) and symptomatic (120 ± 6 days) stages. ALS90 and ALS120 refers to SOD1G93A animals at 90 and 120 days, respectively; WT90 and WT120 correspond to wild-type control animals at the same ages. Statistical differences were determined using t-test considering: * p ≤ 0.05, ** p ≤ 0.01, *** p ≤ 0.001. Data are presented as mean ± SD.

Additionally, the plasma concentrations of the bile acid precursors 27-OH, 3β-HCA and 7α-OH were also greater in symptomatic ALS rats (Fig. 2). While 7α-OH is also formed by nonenzymatic free radical oxidation, other oxysterols commonly linked to oxidative stress, such as 7-ketocholesterol and 5β,6β-epoxycholesterol, did not significantly differ between the groups (Fig. S2).

**Figure 2.**
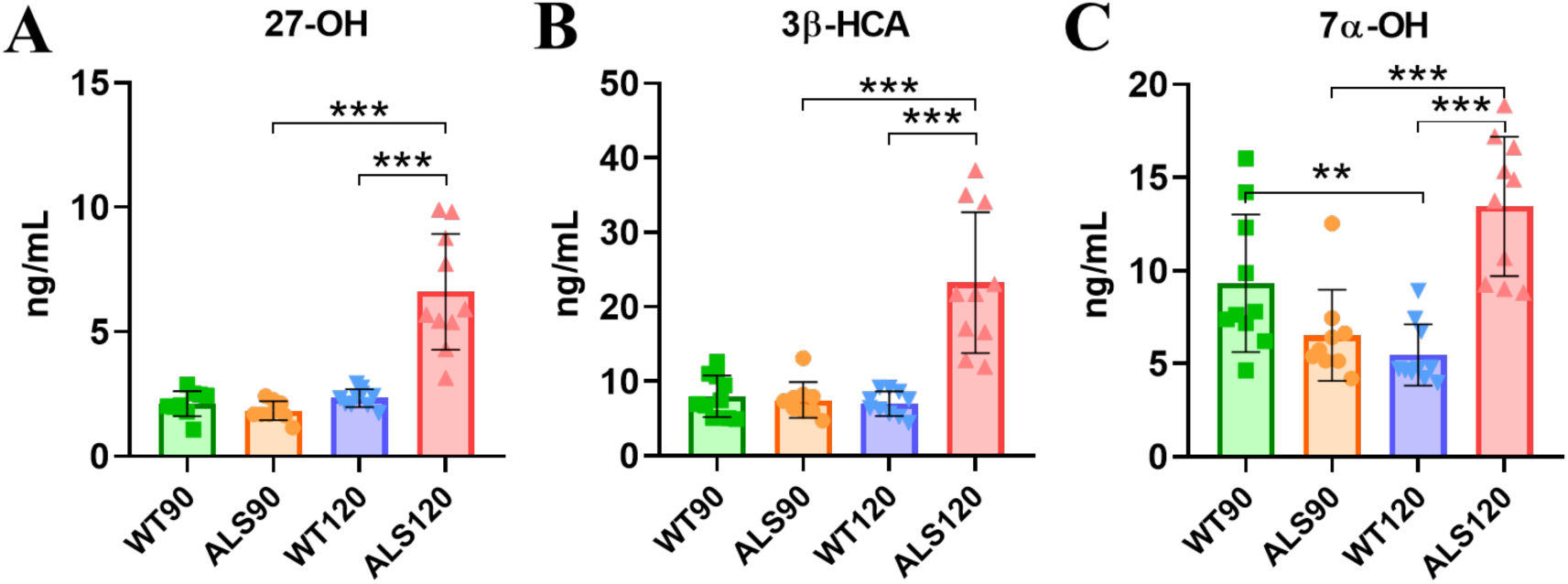
Increased plasma bile acid precursors levels in symptomatic ALS animals. 27-hydroxycholesterol (27-OH, **A**), 3β-hydroxy-5-cholestenoic acid (3β-HCA, **B**) and 7α-OH-cholesterol (7α-OH, **C**) were determined in the blood plasma of asymptomatic and symptomatic animals and age-matched controls at the asymptomatic (89 ± 3 days) and symptomatic (120 ± 6 days) stages. ALS90 and ALS120 refers to SOD1G93A animals at 90 and 120 days, respectively; WT90 and WT120 correspond to wild-type control animals at the same ages. Statistical differences were calculated using t-test: * p ≤ 0.05, ** p ≤ 0.01, and *** p ≤ 0.001. Data are presented as mean ± SD.

The observed increase in bile acid precursors suggests increased bile acid synthesis during advanced stages of the disease. These findings are consistent with previous reports of elevated bile acid concentrations in the spinal cord and feces of end-stage SOD1G93A mice (**Dodge et al., 2021**).

### Spinal cord oxysterol profile reflects neuroinflammation in the ALS model

To further investigate oxysterol alterations in the ALS model, we analyzed the spinal cord. In symptomatic ALS rats, the spinal cord levels of 24(S)-OH were lower than those in age-matched controls (Fig. 3). This reduction can be related to motor neuron loss, as cholesterol 24-hydroxylase (CYP46A1), the CNS-specific and rate-limiting enzyme responsible for conversion of cholesterol to 24(S)-OH, is expressed mainly in neurons [5] or as a result of increased BSCB permeability as the disease progresses. An age-dependent increase in spinal cord 24(S)-OH levels was also observed when 90-day-old and 120-day-old rats were compared. Additionally, 27-OH, an oxysterol mainly synthesized in the liver, was increased in the spinal cord of the ALS 120-day-old animals compared with that in the spinal cord of the asymptomatic 90-day-old animals (Fig. 3), likely reflecting its increase in the blood plasma (Fig. 2A).

**Figure 3.**
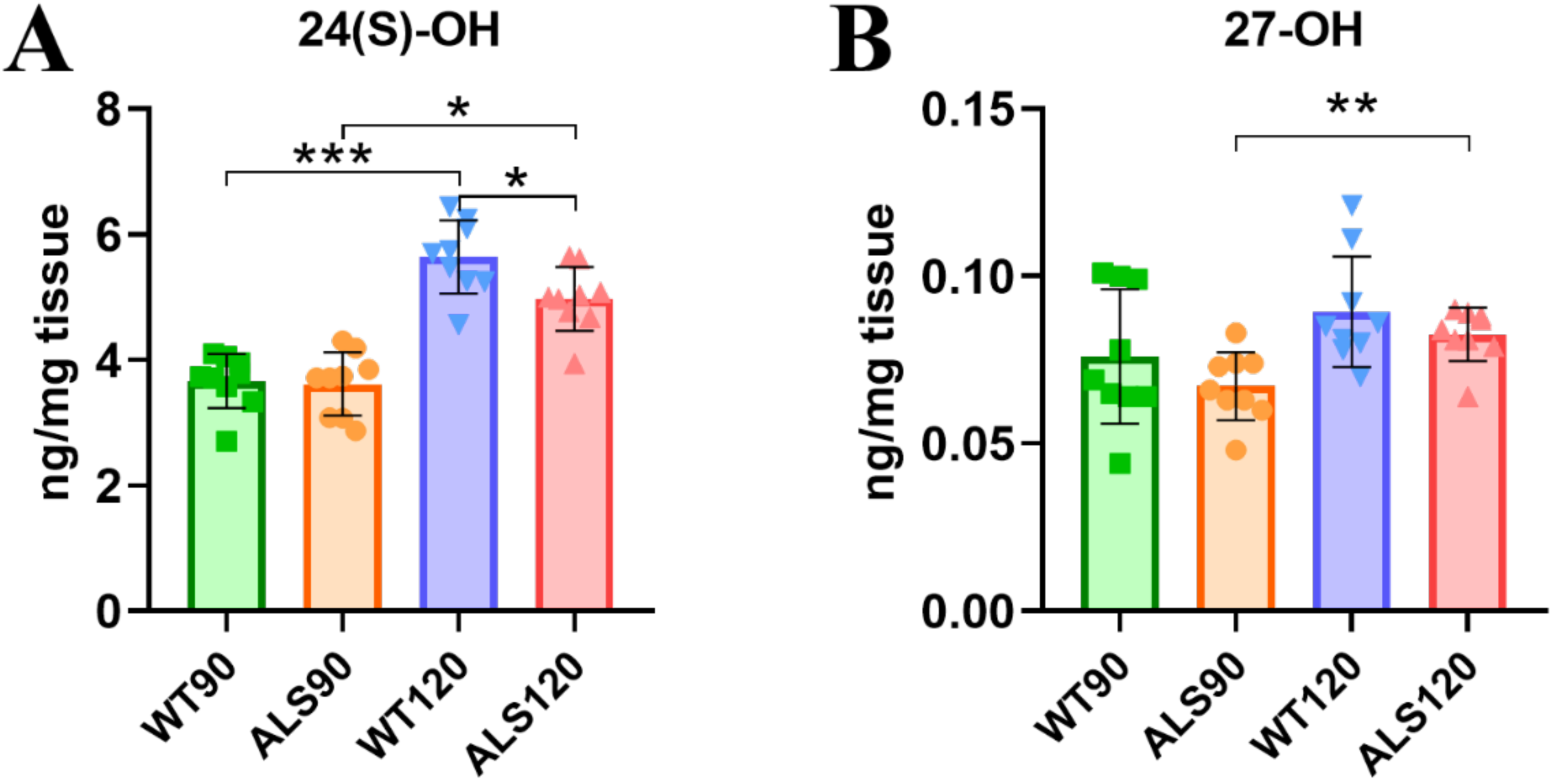
Decreased 24(S)-OH concentration in the spinal cord of ALS symptomatic rats compared to age-matched WT controls (**A**). Increased 27-OH concentration in the spinal cord of symptomatic ALS rats (**B**). ALS90 and ALS120 refers to SOD1G93A animals at 90 and 120 days, respectively; WT90 and WT120 correspond to wild-type control animals at the same ages. Statistical differences were determined using t-test: * p ≤ 0.05, ** p ≤ 0.01, and *** p ≤ 0.001. Data are presented as mean ± SD.

With respect to the neuroinflammatory process, we detected 2–3-fold greater levels of 25-OH in the spinal cord of the symptomatic SOD1G93A rat group than in those of the asymptomatic 90-day-old and aged matched control groups (Fig. 4). Notably, 25-OH was also significantly elevated in the ALS rats at 90 days, indicating that this increase occurred even before symptom onset in the ALS model. These findings support the hypothesis that neuroinflammation may underlie neurodegenerative mechanisms.

**Figure 4.**
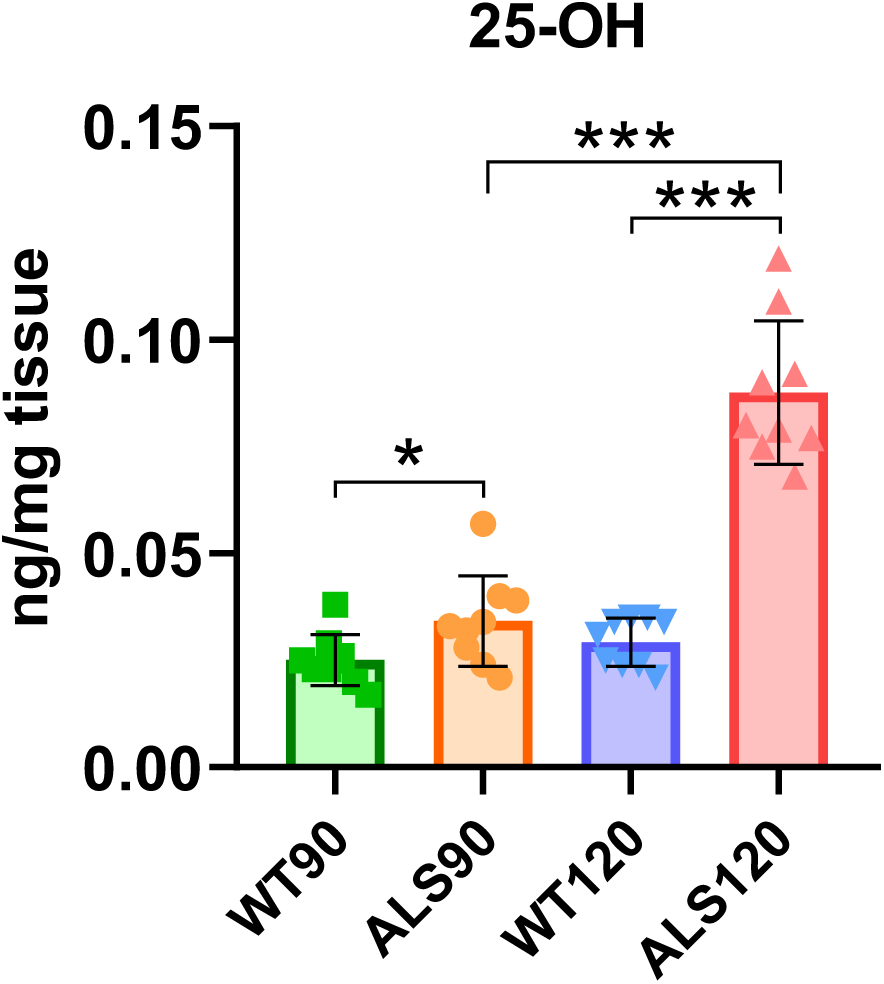
Increased concentrations of 25-OH in the spinal cord of symptomatic (ALS120) and asymptomatic (ALS90) animals. The figure shows a trend toward an increase in 25-OH in the spinal cord during the course of the disease. ALS90 and ALS120 refers to SOD1G93A animals at 90 and 120 days, respectively; WT90 and WT120 correspond to wild-type control animals at the same ages. Statistical differences were calculated using t-test considering: * p ≤ 0.05, ** p ≤ 0.01, *** p ≤ 0.001. Data are presented as mean ± SD.

### Correlation between plasma and spinal cord oxysterols

To investigate the relationship between plasma and spinal cord oxysterols during neurodegeneration, a Pearson correlation was conducted to identify potential links among the altered compounds. In the blood plasma, a strong positive correlation between 25-OH and 24(S)-OH was found (Fig. 5) for symptomatic rats (ALS 120 days), which may be the consequence of an inflammatory condition leading to increased permeability of the BSCB.

**Figure 5.**
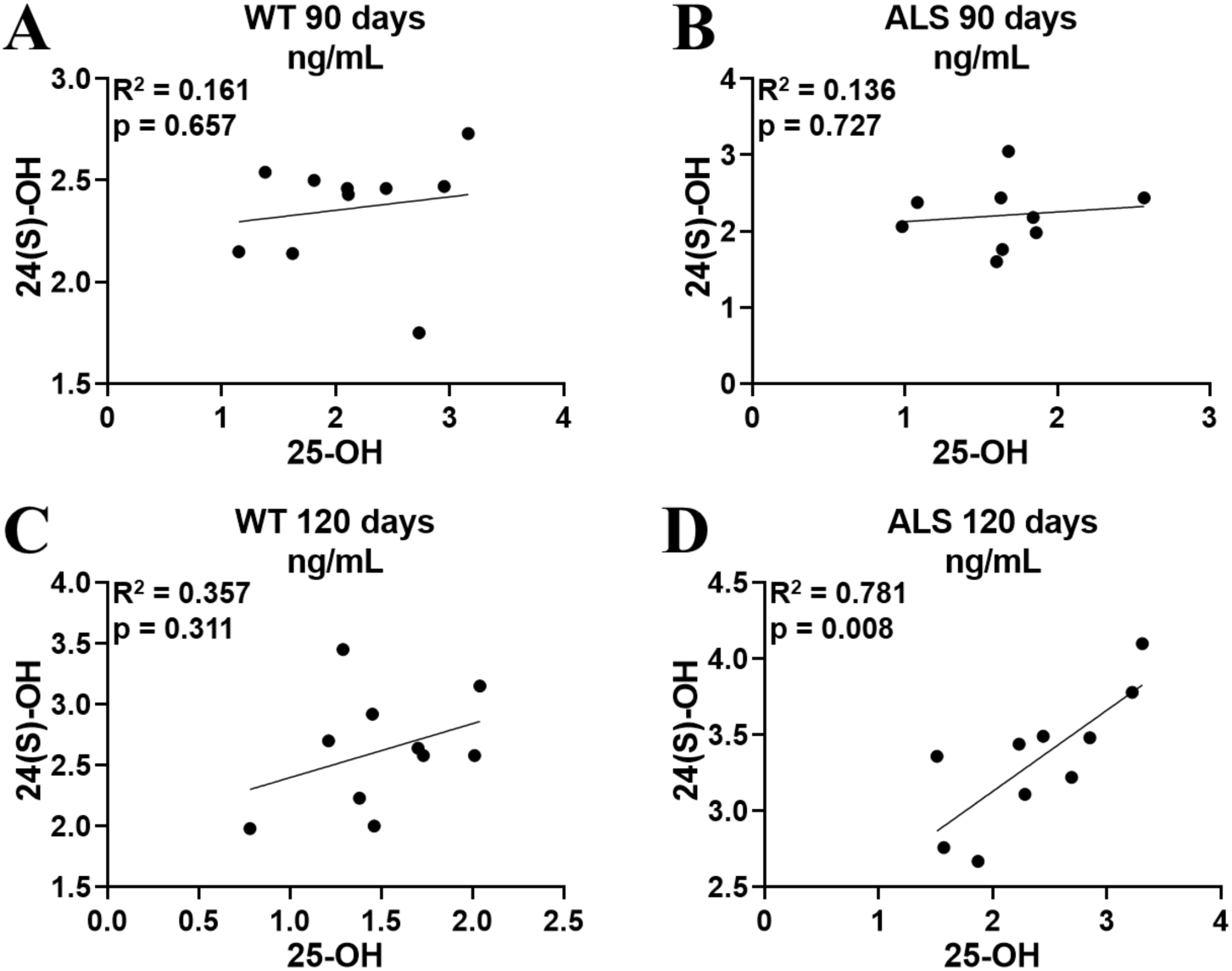
Pearson’s correlation between 24(S)-OH and 25-OH in the plasma of the different groups and ages. A statistically significant correlation was found only in symptomatic SOD1G93A animals (ALS 120 days).

In contrast, in the central nervous system, a strong negative correlation was found between 25-OH and 24(S)-OH (Fig. 6), suggesting that 25-OH may be involved in neuronal loss, consequently reducing the enzyme responsible for 24(S)-OH biosynthesis. Notably, this correlation was observed only in the symptomatic ALS 120-day group, emphasizing the importance of understanding oxysterol alterations in neurodegeneration.

**Figure 6.**
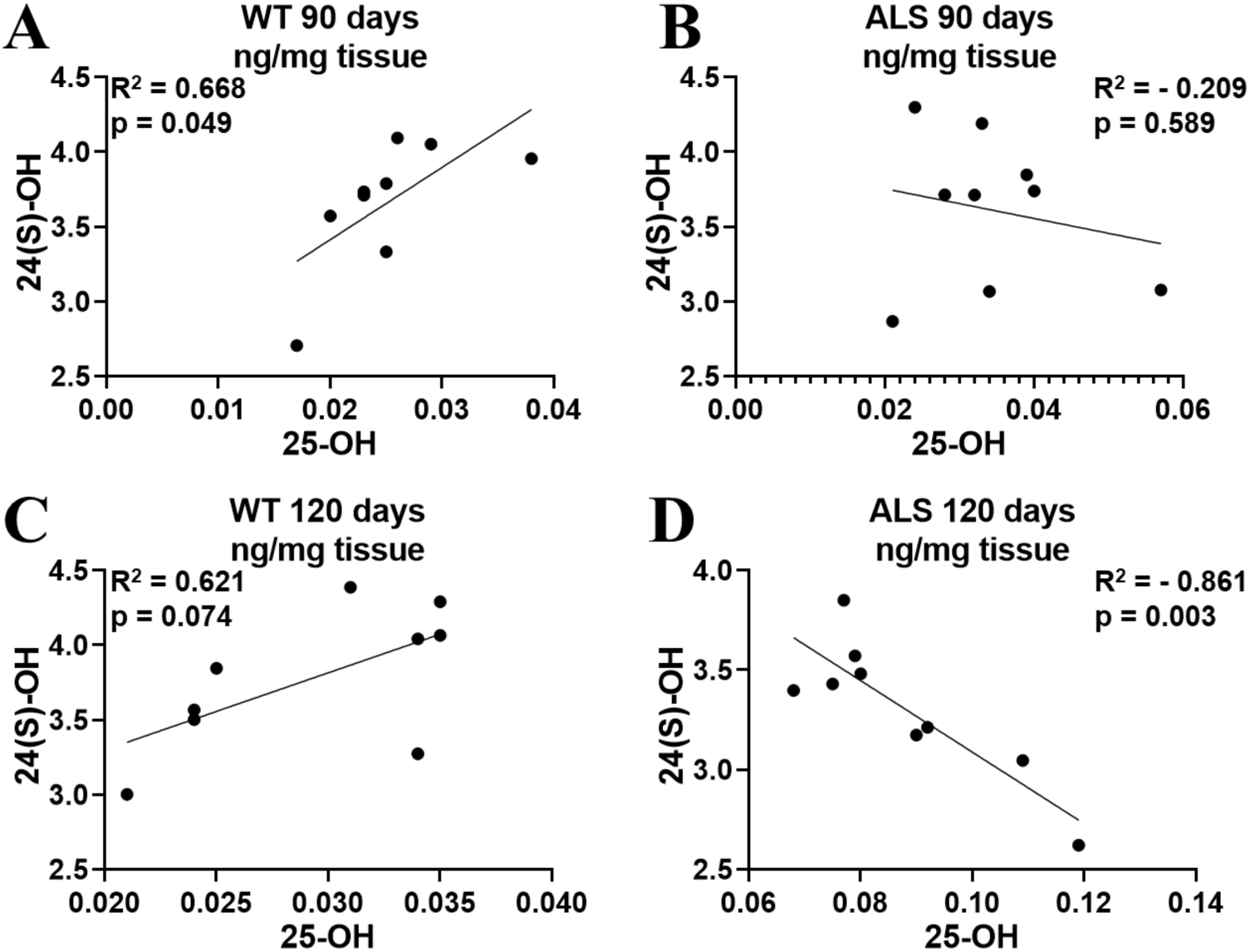
Pearson’s correlation between 24(S)-OH and 25-OH in the spinal cord of the different groups and ages. In symptomatic SOD1G93A animals, the correlation was negative, reflecting the neurodegenerative state, whereas in both control groups, there was a trend for a positive correlation.

Curiously, in the spinal cord of the 90-day control group, an unexpected positive correlation between 25-OH and 24(S)-OH was detected (Fig. 6), possibly reflecting a yet unknown physiological regulatory role of 25-OH produced in situ in CNS cholesterol metabolism. In the 120-day control group, the results did not reach statistical significance but tended toward the positive correlation observed in younger healthy animals (Fig. 6).

### Quantification of plasma LBP in the SOD1G93A rat model

To explore the link between increased intestinal permeability, inflammation, and disease progression, we quantified plasma LBP in SOD1G93A rats via ELISA. LBP is an important protein in the pathway of LPS recognition by TLR4, where it interacts with LPS micelles and facilitates the transfer of LPS monomers to CD14 [64, 65]. LBP is also a marker of intestinal permeability [57, 58] and is used to monitor LPS translocation into the circulation [66]. Recently, increased concentrations of LBP were identified in the plasma of ALS patients, with a positive correlation between LBP concentrations and disease severity [46, 47]. However, no data on LBP quantification in ALS models have been published to date.

The results revealed a significant increase in plasma LBP levels during the asymptomatic stage of the disease. SOD1G93A rats at 90 days presented higher LBP levels than all the other groups, while no significant differences were observed among the remaining groups (Fig. 7). This finding is consistent with previous studies showing increased intestinal permeability in the SOD1G93A mouse model [67], even at asymptomatic stages [45].

**Figure 7.**
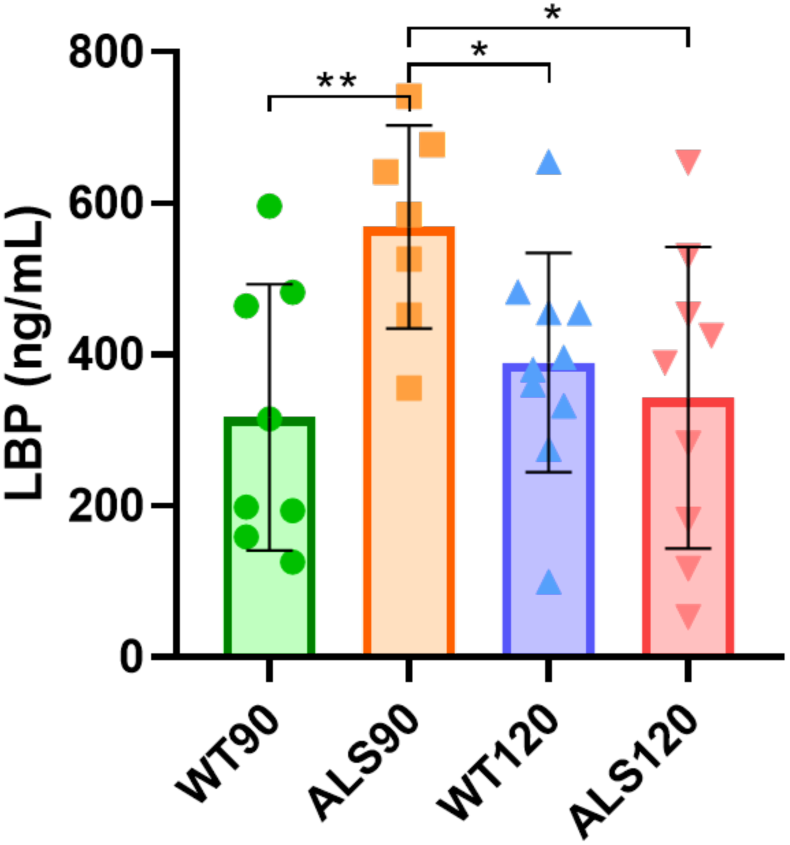
Increased LBP concentrations in the plasma of asymptomatic ALS rats. ALS90 and ALS120 refers to SOD1G93A animals at 90 and 120 days, respectively; WT90 and WT120 correspond to wild-type control animals at the same ages. Statistical differences were calculated using t-test: * p ≤ 0.05, ** p ≤ 0.01, *** p ≤ 0.001. Data are presented as mean ± SD.

Further analysis via Person’s correlation revealed a positive correlation between plasma LBP and 25-OH levels in the spinal cord of asymptomatic rats (ALS 90 days) (Fig. 8). These findings suggest that increased circulating LPS observed at early stages of the disease contributes to the neuroinflammatory process.

**Figure 8.**
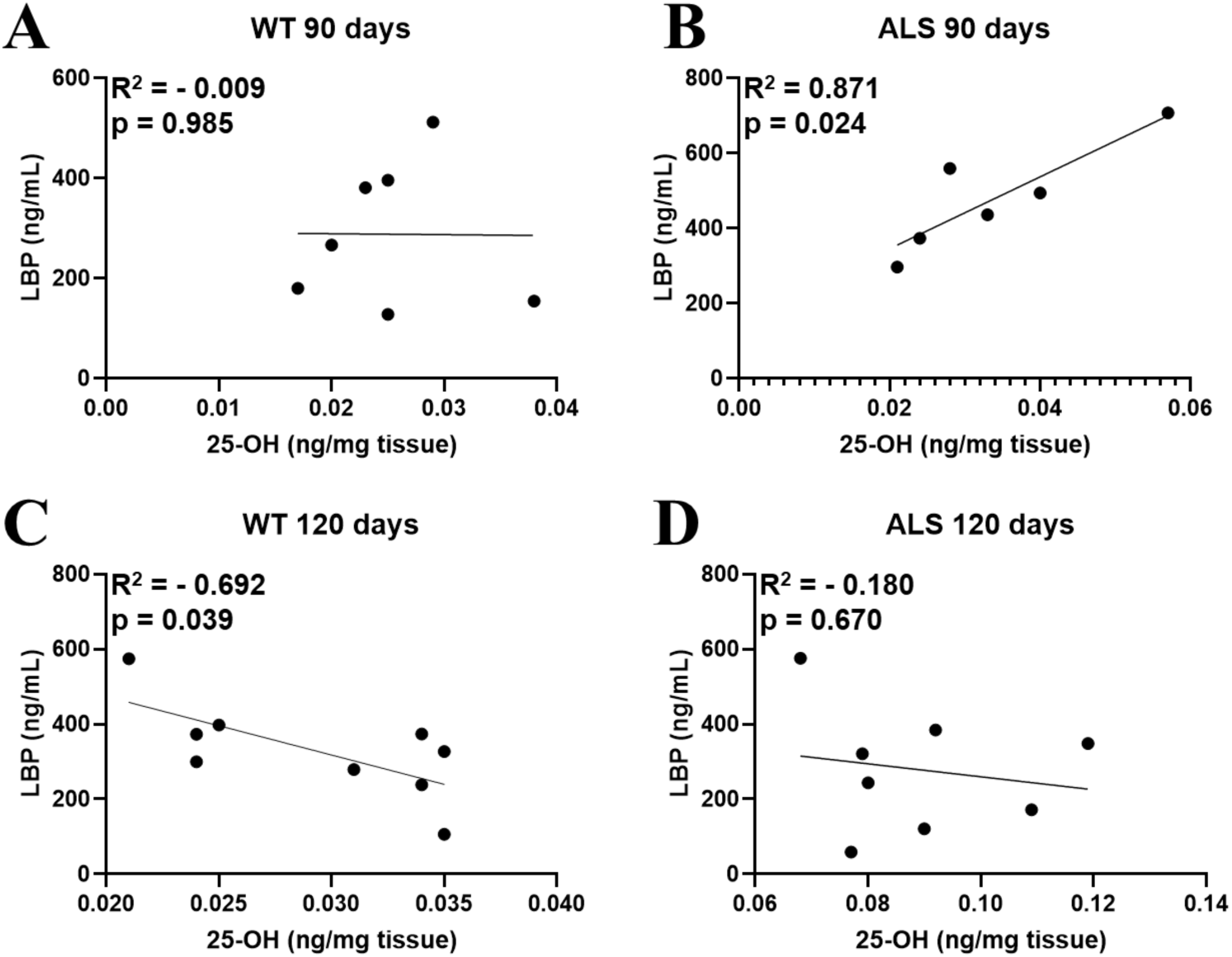
Pearson’s correlation between the plasma concentration of LBP and the levels of 25-OH in the spinal cord of the four groups analyzed. A positive correlation was observed in the ALS 90-day group, which interestingly contained more LBP.

## DISCUSSION

The central nervous system (CNS) is highly enriched in cholesterol, which is synthesized primarily *in situ* by resident cells. Cholesterol homeostasis in CNS is tightly regulated, with oxysterols playing key roles in sterol biosynthesis regulation [16, 17], cholesterol efflux and intercellular transport [68–70] and, immune cell migration [71–73].

Growing evidence indicates that cholesterol and oxysterol metabolism are disrupted during ALS progression [34–38, 74–77]. To investigate this, we quantified oxysterol levels in the blood plasma and spinal cord of SOD1G93A ALS rats by using high-resolution LC-MS analysis. In symptomatic rats, plasma oxysterol levels were significantly increased, particularly the bile acid precursors — 7α-OH, 27-OH, and 3β-HCA — indicating activation of both classical and alternative (acidic) bile acid biosynthesis pathways [10] (Scheme 1). These findings align with previous reports showing increased bile acid levels in the spinal cord and feces of end-stage SOD1G93A mice [38].

Bile acids are emerging as key modulators of metabolic and immune responses. They act on various receptors, including G protein-coupled bile acid receptor 1 (GPBAR1, also known as TGR5), whose activation reduce pro-inflammatory cytokines (IL-12 and TNF-α) in LPS-stimulated macrophages [78]. GPBAR1/TGR5 is also expressed in microglia, where its activation by tauroursodeoxycholic acid (TUDCA) suppresses the expression of pro-inflammatory markers and promotes an anti-inflammatory profile [79]. Clinical trials suggest that TUDCA slows ALS progression, likely by reducing neuroinflammation and preventing protein misfolding and oxidative stress [79, 80].

Gut dysbiosis and intestinal permeability contribute to ALS pathophysiology in both animal models [45, 67, 81, 82] and patients [83–87]. To evaluate alterations in intestinal permeability, we measured plasma level of LPS-binding protein (LBP), an acute phase protein mainly synthesized in the liver [58]. LBP binds to the lipid A moiety of LPS which can derive from translocation from the intestine. It has been shown that high LBP concentrations can inhibit LPS binding to monocytes and induction of proinflammatory cytokines [88]. Our results showed elevated circulating LBP levels in presymptomatic rats, indicating LPS systemic translocation at early disease stage likely due to a disrupted intestinal barrier. However, as the disease progressed, LBP levels declined in symptomatic rats, potentially reflecting a failure to suppress inflammation.

Disrupted intestinal barrier, including reduced levels of tight junction protein ZO-1 in ALS mouse models [45, 67] and elevated plasma LBP levels in ALS patients [46, 47], further support the link between intestinal permeability and systemic inflammation. Our study revealed a positive correlation between increased circulating LBP levels in presymptomatic rats and elevated 25-hydroxycholesterol (25-OH) levels in the spinal cord, supporting a link between gut permeability, LPS and neuroinflammation in the CNS.

LPS lipid A structures from various bacterial strains can act as either TLR4 agonists or antagonists, influencing pro-inflammatory signaling [89, 90]. Thus identifying the bacterial sources and LPS structures involved in ALS could provide valuable insights. Additionally, AOAH, an enzyme that deacylates inflammatory LPS to its nonstimulatory forms [91], relies on LBP for efficient activity [92]. Interestingly, assessment of transcriptomic data (R Shiny App) from ALS patient spinal cord samples revealed increased expression of AOAH, CD14, and CH25H, further linking LPS and alterations in oxysterol metabolism to neuroinflammation [93].

Emerging evidence indicates that 25-OH and its dihydroxylated derivative 7α,25-dihydroxycholesterol plays a key role in neuroinflammation [72, 73]. Synthesized by macrophages and microglia upon LPS stimulation [94], 25-OH is emerging as important regulators of immune responses and are implicated in CNS homeostasis and neurodegenerative diseases [95–97]. Notably, in the ALS rat model spinal cord 25-OH levels were elevated in both presymptomatic and symptomatic stages, with a two-fold increase in symptomatic animals. Its early elevation at asymptomatic stage suggests that immune activation precedes symptom onset and intensifies as the disease progresses.

In plasma, 25-OH correlated positively with the increase in 24(S)-OH levels. The observed increase in plasma levels of 24(S)-OH, a CNS-derived oxysterol primarily synthesized by neurons [6, 12], in symptomatic rats may reflect blood‒ brain barrier (BBB) and blood‒spinal cord barrier (BSCB) disruption [39–44]. As these barriers become compromised in ALS, erythrocytes [44], immune cells [56] and other blood components, including oxysterols, can infiltrate the CNS (Scheme 1). Elevated blood-derived 27-OH levels in the spinal cord of symptomatic rats also support the idea that barrier dysfunction is associated with disease progression.

**Scheme 1.**
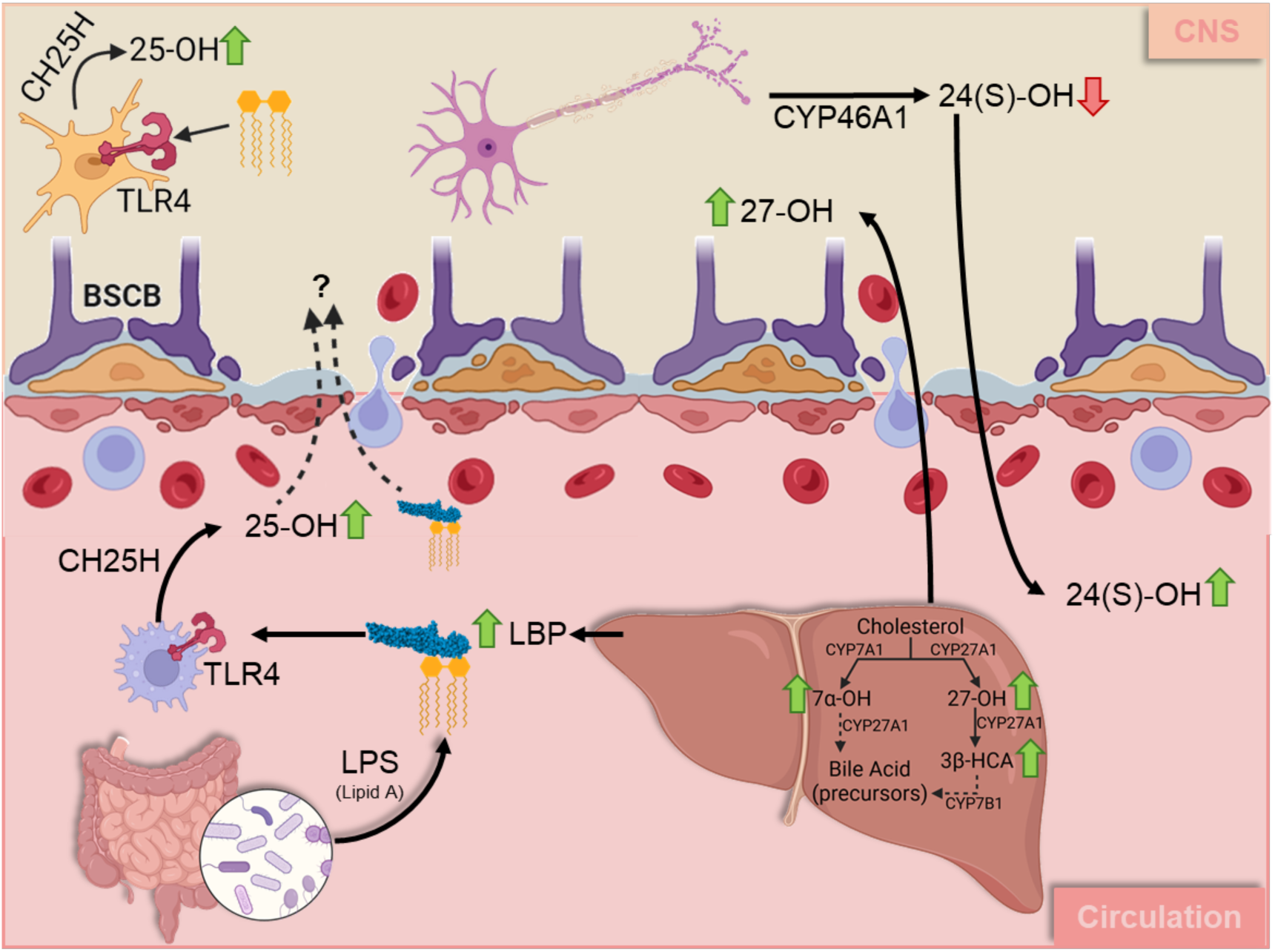
Dysrupted oxysterol metabolism in ALS. Lower panel left: Increased LBP levels indicate enhanced intestinal permeability and subsequent LPS translocation into systemic circulation. LPS stimulation of macrophages activates CH25H, leading to increased 25-hydroxycholesterol (25-OH) production at the presymptomatic stage, which suggests an early neuroinflammatory process. A positive correlation between plasma levels of 24(S)-OH and 25-OH in symptomatic rats further support a link between immune activation and increased BBB/BSCB permeability. Lower panel right: Cholesterol metabolism in the liver shows increased levels of bile acid precursors 7α-OH, 27-OH, and 3β-HCA, reflecting the activation of both classical and alternative (acidic) bile acid biosynthesis pathways. Upper panel: In the spinal cord, microglia activation results in increased 25-OH production (left) accompanied by reduced levels of 24(S)-OH. Since 24(S)-OH is produced by CYP46A1, primarily located in neurons, its decrease is likely related to motor neuron loss during disease progression (right).

In the spinal cord, 25-OH levels correlated negatively with 24(S)-OH, suggesting that microglial activation, characterized by increased 25-OH production [98, 99], contributes to neuronal loss, leading to reduction in 24(S)-OH levels. 25-OH has been shown to induce apoptosis in NSC34 neuronal cell by activating the GSK-3β/ LXR pathway [35]. In a recent study Urano et al. demonstrated that 25-OH induces ferroptosis by downregulating GPX4 expression [97]. These findings highlight the multifaceted neurotoxic role of 25-OH in ALS pathology, linking microglial activation to neuronal loss through both apoptotic and ferroptotic mechanisms.

Microglial and neuroimmune activation are hallmarks of ALS [48–52], yet their precise mechanisms remains uclear. Emerging evidence suggests that oxysterols, are potent ligands of the immune receptor G-protein coupled receptor Epstein-Barr virus-induced gene 2 (EBI2), also known as GPR183, functioning as chemoattractants of immune cells [71–73]. Interestingly, GPR183 is also expressed in microglia and astrocyte membranes [100]. Moreover the enzymes Ch25h and CYP7B1, which synthesize its most potent ligand, 7α,25-diOH, are increased in LPS-stimulated glial cells [98, 99, 101], emphasizing the role of oxysterols in neuroimmune signaling.

## Conclusion

This study provides new insights into oxysterol dysregulations in ALS, demonstrating an early increase in 25-OH production from presymptomatic stages, with further elevations as disease progresses. Additionally, elevated LBP levels in presymptomatic SOD1G93A rats, suggest a link between early inflammatory activation and intestinal permeability dysfunction. The correlation between LBP and spinal cord 25-OH levels further supports a connection between gut permeability and neuroinflammation in ALS. Further studies are required to confirm these findings and explore the mechanisms underlying the interplay between intestinal barrier dysfunction, oxysterol metabolism and neurodegeneration.

## Declarations

### Ethics approval

All experimental procedures were conducted in accordance with the Ethics Committee on Animal Use of the University of São Paulo (CEUA 41/2016).

### Consent for publication

Not applicable.

### Availability of data and materials

The datasets generated during and/or analyzed during the current study are available from the corresponding author upon reasonable request.

### Authors’ contributions

SM and RSL conceived and designed the study. RSL, RSS and IFLT conducted the experiments and performed the analyses with guidance from SM and AAS. SM and RSL wrote the manuscript and analyzed the data. ABCF, AI, RLF, LSD and IFDP conducted the animal experiments. SM, MHGM and AS critically revised the manuscript. All the authors read and approved the final manuscript.

## Acknowledgments

Not applicable.

