## Supplementary Tables and Figures for "Oxysterol Alterations in SOD1G93A ALS Rats: 25-Hydroxycholesterol and LPS-Binding Protein in Disease Progression"

### Spinal cord 25-OH alterations in $SOD1^{G93A}$ rats correlate with plasma LPS-binding protein

Supplementary Table1: Standards in the oxysterol mix solution and deuterated IS used in samples

Supplementary Table2: Calculated parameters of the oxysterols calibration curves

Figure S1: Chromatogram of oxysterols and sterol standards with their respective monitored m/z.

Figure S2: Levels of oxysterols (**A**) and sterols (**B**) in the plasma of SOD1G93A and control groups at 90 and 120 days old.

Figure S3: Levels of oxysterols (**A**) and sterols (**B**) in the spinal cord of SOD1G93A and control groups at 90 and 120 days old.

**Supplementary Table 1**. Standards in the oxysterol mix solution and deuterated IS used in samples

| **Sterol standards** | | | **Source** | |
|---|---|---|---|---|
| **Mix of oxysterols** | **Abbreviation** | **Concentration** | **Code** | **Company** |
| 7-ketocholesterol | 7-Keto | 2 µM | 700015P | Avanti |
| 7α-hydroxycholesterol | 7α-OH | 2 µM | 700034P | Avanti |
| 5β,6β-epoxycholesterol | 5β,6β-EC | 2 µM | 700033P | Avanti |
| 25-hydroxycholesterol | 25-OH | 2 µM | 700019P | Avanti |
| 24(S)-hydroxycholesterol | 24(S)-OH | 2 µM | 700061 | Avanti |
| 27-hydroxycholesterol | 27-OH | 2 µM | 700021 | Avanti |
| 4β-hydroxycholesterol | 4β-OH | 2 µM | 19518 | Cayman |
| 22(R)-hydroxycholesterol | 22(R)-OH | 2 µM | 89355 | Cayman |
| 3β-hydroxy-5-cholestenoic acid | 3β-HCA | 2 µM | 21859 | Cayman |
| 5α-hydroxycholesterol | 5α-OH | 2 µM | Synthesized | |
| 6β-hydroxycholesterol | 6β-OH | 2 µM | | |
| 3β-hydroxy-5-oxo-5, 6-secocholestan-6-al | Csec | 2 µM | | |
| 3β-hydroxy-5β-hydroxy-B-norcholestane-6β-carboxyaldehyde | ChAld | 2 µM | | |
| **Internal Standards** | **Abbrevation** | **Concentration** | **Code** | **Company** |
| d6 24-hydroxycholesterol | d6 24-OH | 0.5 µM | LM-4110 | Avanti |
| d7 7-ketocholesterol | d7 7-keto | 0.5 µM | LM-4107 | Avanti |
| d7 7α-hydroxycholesterol | d7 7α-OH | 0.5 µM | LM-4103 | Avanti |
| d6 Cholesterol | d6 Chol | 2.5 mM | 488577 | ISOTEC |

**Supplementary Table 2 - Calculated parameters of the oxysterols calibration curves**

| Oxysterol | IS | $R^2$ | Average CV % (low - high) | Accuracy % (low - high) | LOQ *on-column* |
|---|---|---|---|---|---|
| **3β-HCA** | | 0.9955 | 4.4 (1 - 12) | 88 - 107 | 13.2 pg (32.8 fmol) |
| **22(R)-OH** | | 0.9940 | 3.6 (2 - 8) | 85 - 110 | 5.28 pg (13.1 fmol) |
| **25-OH** | d6 24-OH | 0.9944 | 3.6 (1 - 8) | 85 - 106 | 5.28 pg (13.1 fmol) |
| **24(S)-OH** | | 0.9936 | 4 (1 - 9) | 85 - 108 | 5.28 pg (13.1 fmol) |
| **27-OH** | | 0.9941 | 2.9 (1 - 7) | 85 - 110 | 5.28 pg (13.1 fmol) |
| **CSec** | d7 7-Keto | 0.9833 | 7.3 (3 - 11) | 95 - 113 | 33.00 pg (82 fmol) |
| **7α-OH** | d7 7α-OH | 0.9921 | 6.7 (3 - 12) | 85 - 109 | 5.28 pg (13.1 fmol) |
| **7-Keto** | d7 7-Keto | 0.9901 | 2.6 (1 - 4) | 99 - 109 | 82.48 pg (204.8 fmol) |
| **ChAld** | | 0.9921 | 7.6 (1 - 17) | 87 - 110 | 5.28 pg (13.1 fmol) |
| **5α-OH** | | 0.9939 | 6.4 (4 - 10) | 88 - 109 | 5.28 pg (13.1 fmol) |
| **5β,6β-EC** | d7 7α-OH | 0.9912 | 5.2 (1 - 11) | 99 - 109 | 82.48 pg (204.8 fmol) |
| **6β-OH** | | 0.9872 | 8.5 (3 - 13) | 85 - 112 | 33.00 pg (82 fmol) |

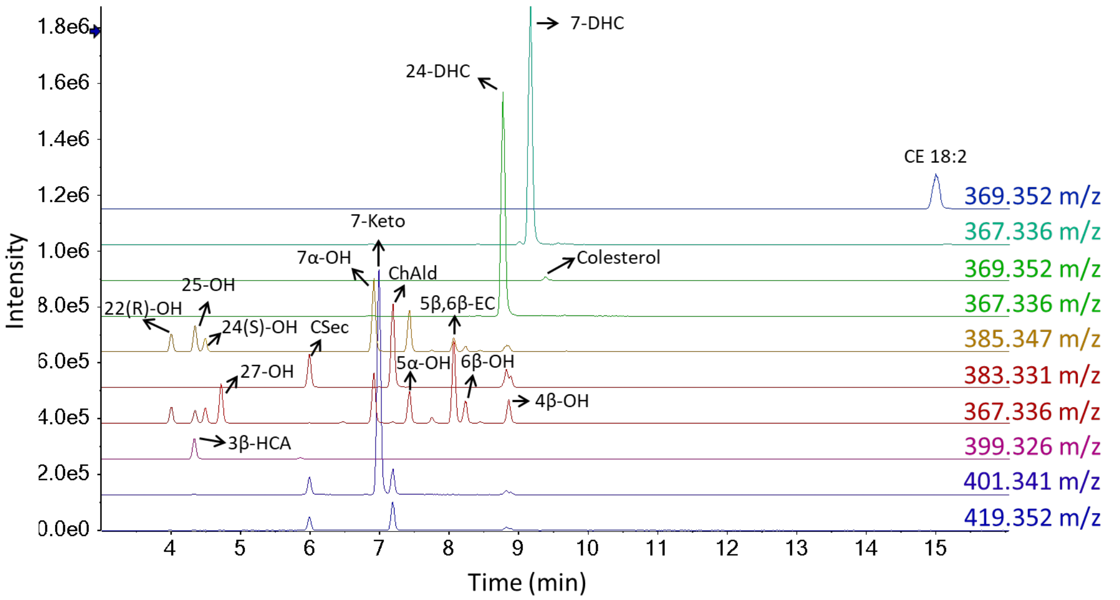

**Figure S1** – Chromatogram of oxysterols and sterol standards with their respective monitored m/z.

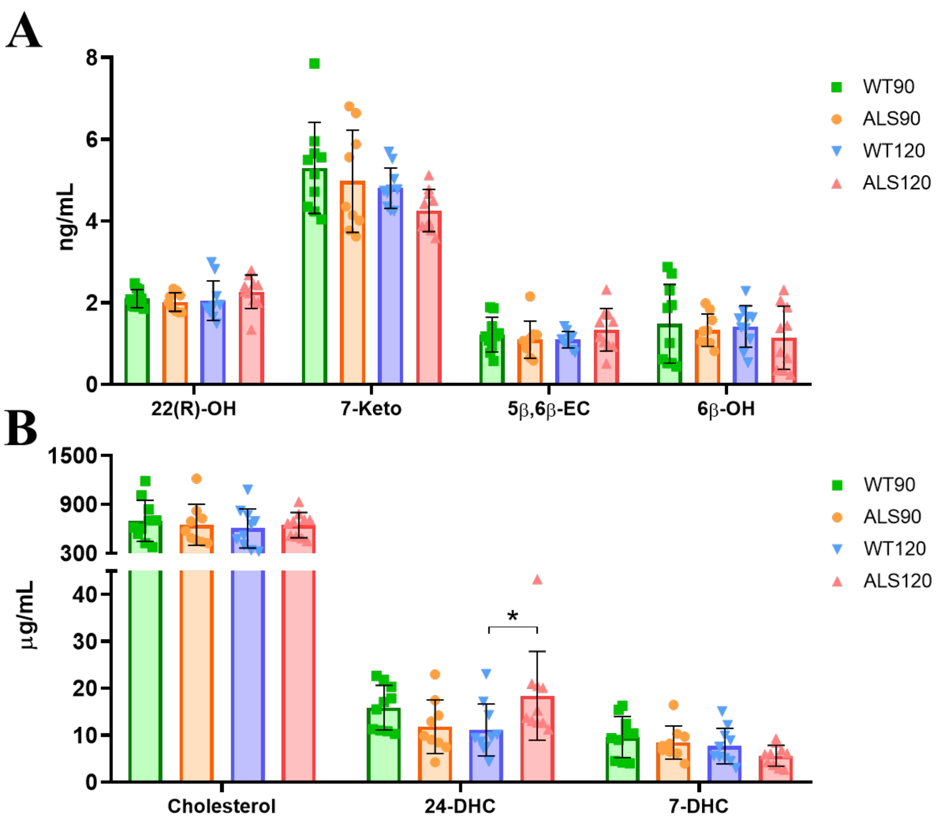

**Figure S2** – Levels of oxysterols (**A**) and sterols (**B**) in the plasma of SOD1G93A and control groups at 90 and 120 days old. Sterols were semi-quantified based on d6-Cholesterol. Statistical differences were calculated using t-test considering: * $p \leq 0.05$, ** $p \leq 0.01$, *** $p \leq 0.001$. The graph shows the mean±SD.

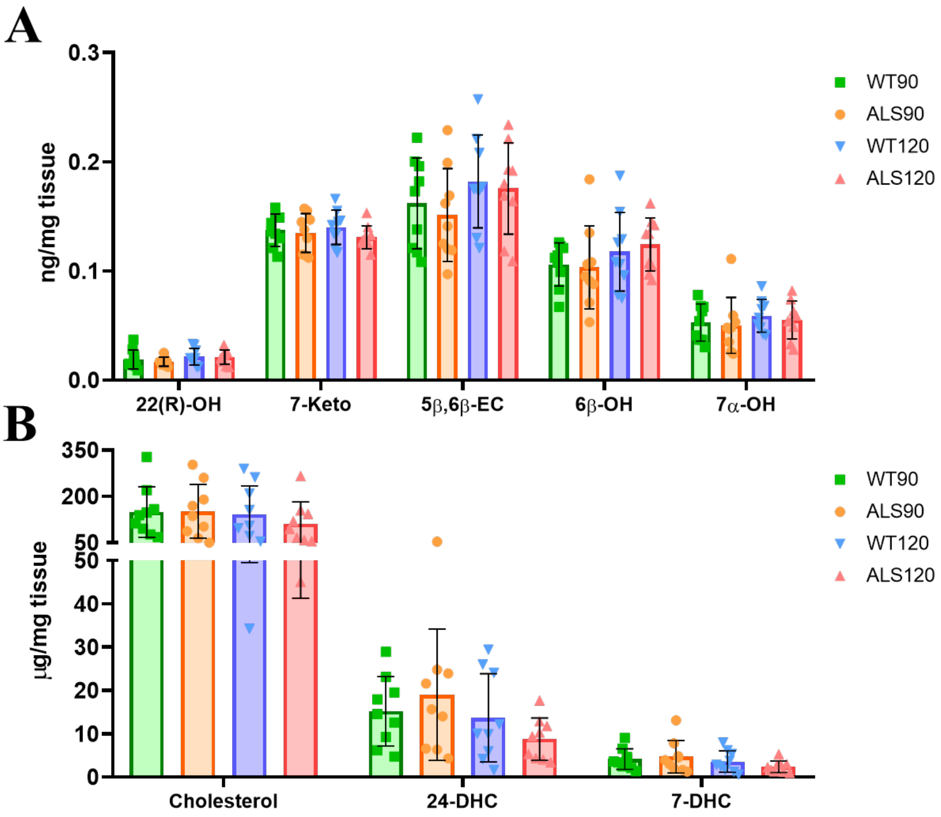

**Figure S3** – Levels of oxysterols (**A**) and sterols (**B**) in the spinal cord of SOD1G93A and control groups at 90 and 120 days old. Sterols were semi-quantified based on d6-Cholesterol. Statistical differences were calculated using t-test considering: * $p \leq 0.05$, ** $p \leq 0.01$, *** $p \leq 0.001$. The graph shows the mean±SD.
